# Why human papillomavirus acute infections matter

**DOI:** 10.1101/144576

**Authors:** Samuel Alizon, Carmen Lía Murall, Ignacio G. Bravo

**Affiliations:** MIVEGEC (UMR CNRS 5290, UR IRD 224, UM), 911 avenue Agropolis, 34394 Montpellier Cedex 5, France

**Keywords:** clearance, persistence, latency, chronic, meta-population, fertility, virome, warts, cancer, evolution, ecology, vaccination

## Abstract

Most infections by human papillomaviruses (HPVs) are ‘acute’, that is non-persistent. Yet, for HPVs, as for many other oncoviruses, there is a striking gap between our detailed understanding of chronic infections and our limited data on the early stages of infection. Here we argue that studying HPV acute infections is necessary and timely. Focusing on early interactions will help explain why certain infections are cleared while others become chronic or latent. From a molecular perspective, descriptions of immune effectors and pro-inflammatory pathways during the initial stages of infections have the potential to lead to novel treatments or to improved handling algorithms. From a dynamical perspective, adopting concepts from spatial ecology, such as meta-populations or meta-communities, can help explain why HPV acute infections sometimes last for years. Furthermore, cervical cancer screening and vaccines impose novel iatrogenic pressures on HPVs, implying that anticipating any viral evolutionary response remain essential. Finally, hints at the associations between HPV acute infections and fertility deserve further investigation given their high worldwide prevalence. Overall, understanding asymptomatic and benign infections may be instrumental in reducing HPV virulence.

---

The most oncogenic viruses to humans are a group of around 20, closely related, Human papillomavirus (HPV) types. All of them are classified in the alpha-papillomaviruses genus and classically referred to as ‘high risk types’ [1]. HPV-induced cancers typically occur after several years of infection (Figure 1). The importance of viral persistence in the natural history of these cancers has driven most research to focus on chronic infections and to relatively neglect acute infections (*sensu* Virgin *et alii* [2]). For instance, when studying the duration of HPV infections in young women, infections that clear within two to three years tend to be referred to only indirectly, i.e. without a qualifying adjective [3]. Here we aim at clarifying what acute HPV infections are and to summarize the current understanding about them, in the context of the recent progress in the fight against HPVs. Finally, we identify the main gaps in our knowledge about such acute infections, which, if filled, would have direct implications for preventing, controlling and treating infections by HPVs.

**Figure 1.**
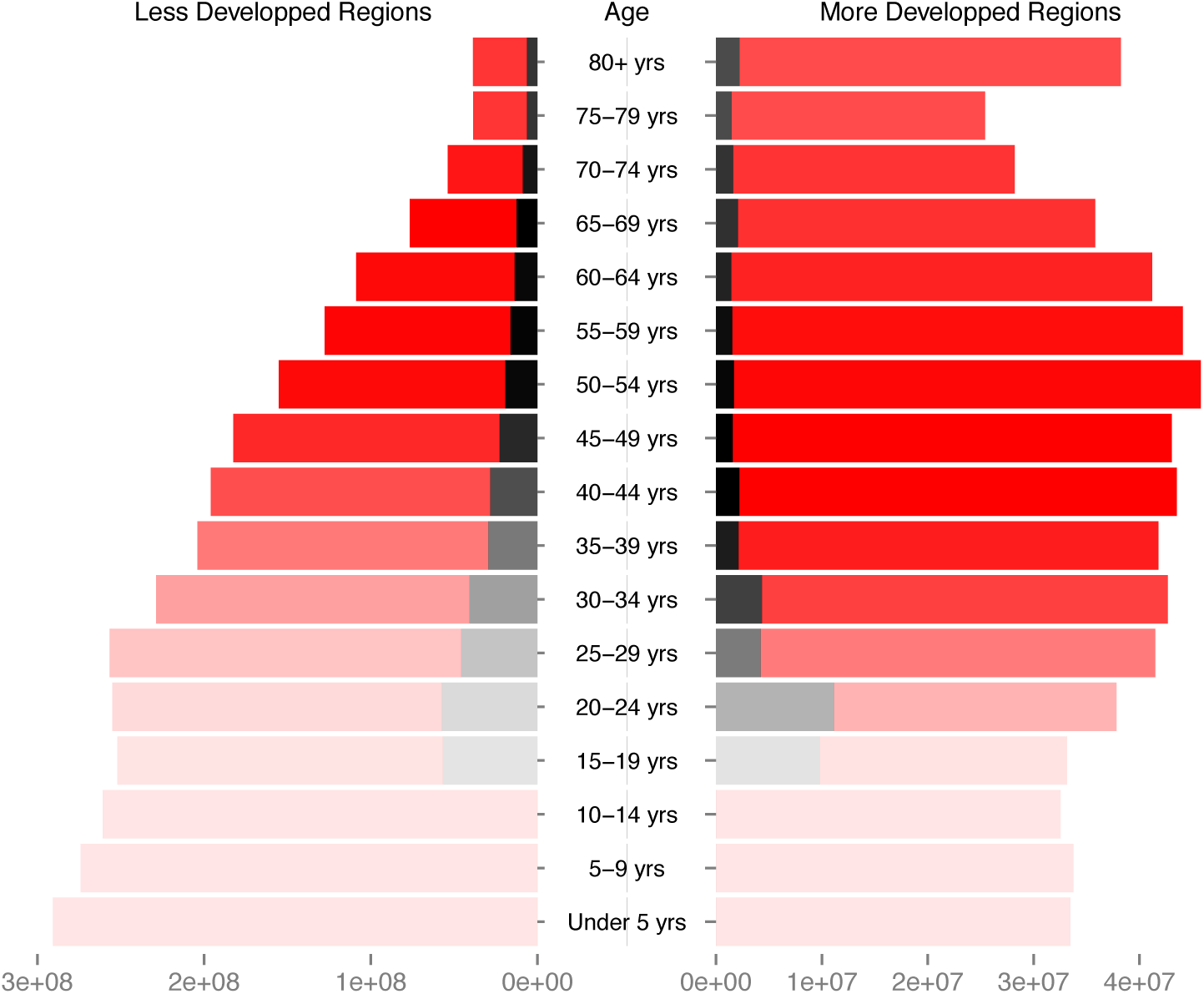
Number of women with asymptomatic cervical HPV infections (black bars) as a function of age class (red bars) and of country development status. Bar color intensity reflects the prevalence of cervical cancer in the corresponding age class (and hence the focus of current research). Data should be read as follows, using the 30-34 years of age as an example: in less developed regions, among the 228 million women in the age class, 40 million (17,8%) display a normal cytology but are actually infected by HPVs, and 30 thousand (0,013%) suffer from cervical cancer, while in more developed regions among the 42 million women in the age class, 4.3 million (10,2%) display a normal cytology but are actually infected by HPVs, and 6 thousand (0,015%) suffer from cervical cancer. Data correspond to HPV prevalence in women with normal cytology, as estimated in the HPV information center (<u>http://hpvcentre.net/</u>, [4]). Overall HPV prevalence values will be actually larger, as they will include women with abnormal cytology. No data were available for HPV prevalence in women below 15 years of age. Demographic data correspond to the UN projections for 2015 (note the shift in the logarithmic scale for the left and the right sides).

## State of the fight against HPVs

Infection-driven cancers are distinctive because they can be fought using the arsenal developed against infectious diseases: identification of risk factors, prevention of transmission and early detection of infected individuals. Identification of risk factors has led to the recognition of a few, closely related oncogenic HPVs as necessary etiologic agents of several cancers [5]. Contagion can now be prevented by the use of safe and effective vaccines targeting the most oncogenic HPVs along with certain non-oncogenic HPVs that cause anogenital warts [6,7]. Screening programs for early detection of (pre)neoplastic lesions caused by HPVs infections have also been successful at decreasing the burden of cervical cancer in rich countries [8]. However, their differential implementation has also increased the inequality between countries [9,10].

In spite of primary and secondary prevention measurements available, HPVs will continue to infect millions of people in the foreseeable future, thereby causing significant morbidity and mortality worldwide [11]. Indeed, vaccine coverage varies widely both within and between countries [6], as does access to screening programs [12]. Beyond socio-economical factors, screening effectiveness is hampered by the fact that certain forms of cancer are more difficult to detect than others. This is the case for glandular forms of cervical cancer compared to the more common squamous carcinoma. They are often overlooked during standard screening procedures and their incidence is increasing [13,14]. Furthermore, cancers induced by HPVs in anatomical locations other than the cervix (e.g. anal [15] or oropharyngeal [16]) are on the rise in many countries, albeit in different populations [5]. These cancers are particularly worrying: either because they are detected once the carcinogenic process is more advanced, as is the case of head and neck cancer [16], or because they affect populations at increased risk, as is the case of anal cancer in HIV-infected men having sex with men [15]. Finally, from an economical perspective, non-carcinogenic HPVs should not be overlooked since the total health care cost linked to treating genital warts can exceed that of treating HPV-induced cancers [17], despite the obvious differences in severity and indirect impact of both diseases.

### HPV acute infections

#### A definition challenge

HPV infections that are not chronic or latent have been referred to as ‘acute’ [18,19], ‘non-persistent’ [20], ‘transient’ [21], or ‘cleared’ [22] infections. Here, following Virgin *et al.* [2], we define acute infections as a non-equilibrium process that results either in infection clearance, host death or chronic infection.

As illustrated in Figure 2, clinical detection patterns may often lead to ambiguities. First, after sexual intercourse with an infected partner, viral genetic material may be detected for several days, even in the absence of an infection. We refer to these as ‘transitory infections’, although ‘transitory detection’ might be more accurate. Second, some infections successfully establish, replicate the viral genome, produce virions, and are eventually cleared (Figure 2). We refer to these as ‘acute infections’. Third, some acute infections only appear to clear, but the viral genome remains in the infected cell without detectable activity [18,24]. We refer to these as ‘latent infections’. The viral genetic material may occasionally be detected during latency. Reactivation of the viral activity in latent infections may occur much later, for instance triggered by immunesupression [25], but often also without any obvious reason. Finally, some acute infections are not cleared and maintain viral activity over time. We refer to these as ‘chronic infections’. Clinically relevant chronic infections may still resolve naturally, sometimes in a matter of years [8,21]. Acute, latent and chronic infections most likely differ in terms of viral activity, e.g. viral and cellular gene expression patterns, effects on cell replication dynamics or induced local immunosuppression [26]. Nevertheless, in the absence of a proper follow-up study design, characterising the stage of an infection remains difficult, as the detection of viral genetic material associated to latent or to chronic infections [2] can bias our estimates about the prevalence of acute infections, as explained below.

**Figure 2.**
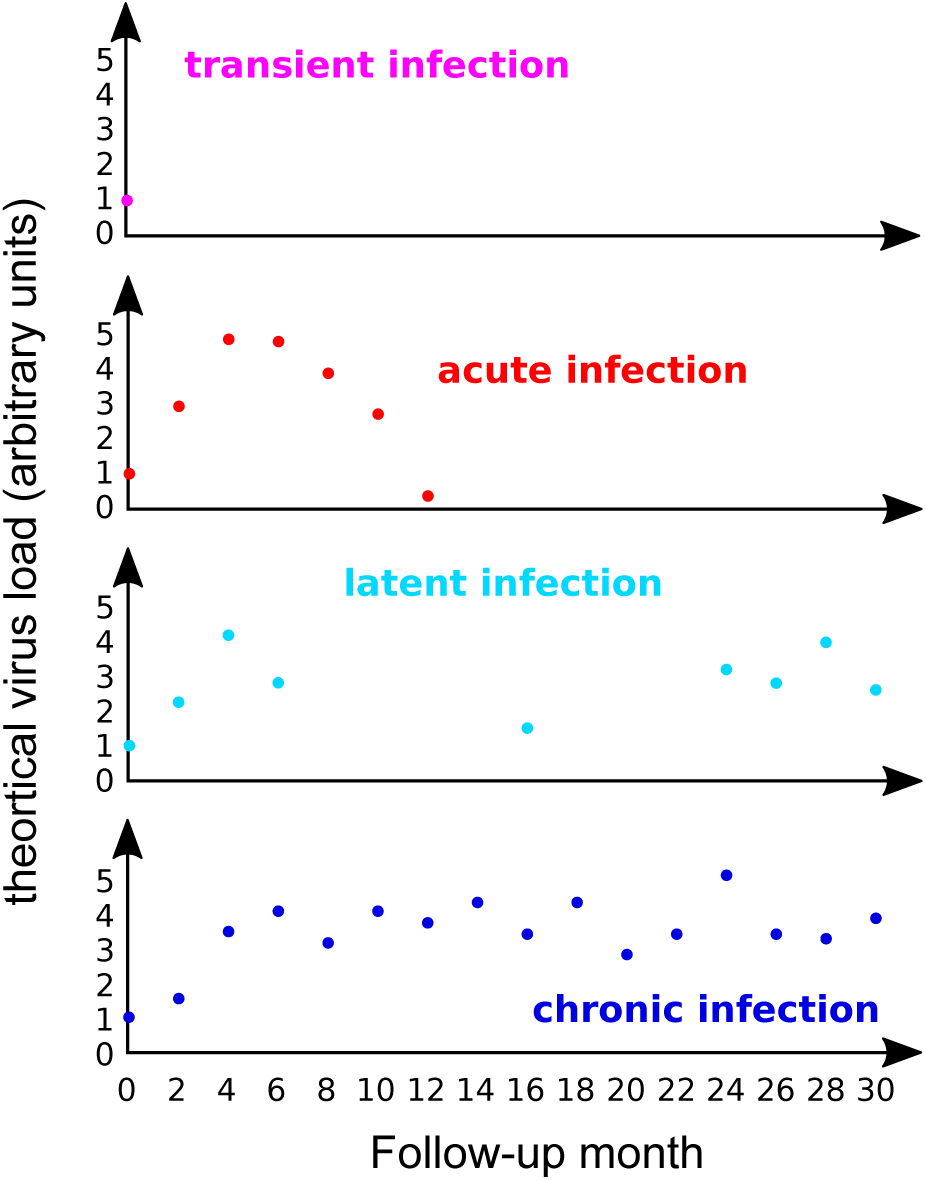
Kinetics of HPV virus load in different genital infections. The panels present made-up data with four imaginary patients monitored every 2 months from the onset of the infection (month 0). Currently most data describe viral load in precancerous lesions and in cancers [23], while very little data focus on how virus loads vary over the course of an acute infection.

One important reason for the blurry line between acute and chronic infections is that HPVs are very diverse in terms of genetics as well as in clinical presentation of the infection. For instance, papillomaviruses colonise the skin and mucosa of virtually all humans from very early in life [27–29] and replicate at very low levels without any apparent clinical or cellular damage. In this respect, humans are continuously being infected (and reinfected) by HPVs, and most humans host a number of latent or chronic infections by HPVs. Certain viral genotypes cause proliferative infections with clinical manifestations such as warts or pre-neoplastic lesions. As we will see below, most of these proliferative lesions resolve naturally albeit in a matter of years, meaning that acute infections by HPVs can be of long duration. Overall, only a detailed understanding of the role of the immune response and clearance, of the nature and permissiveness for latency/chronicity of the cellular targets, and of the timing and repertoire of viral gene expression, will allow us to clarify the frontier between acute and chronic HPV infections.

#### Most infections by HPVs are acute

For most viruses the acute, clinically prominent phases of viral infections, are generally better understood than the chronic ones [2]. Research on oncogenic HPVs is an exception because of its focus on chronicity and on virus-induced cell transformation. Nevertheless, epidemiology data strongly suggest that the vast majority of anogenital infections by oncogenic HPVs never become chronic [8]. In women, the incidence of novel anogenital infections by oncogenic HPVs decreases with age, while persistence increases with age [30,31]. In men, this risk for novel infections is stable with age [32]. For infections in the female genital tract, the prevalence is U-shaped with age [4,33]. For both, men and women, young adults exhibit the highest prevalence, which often rises above 25% (black bars in Figure 1). HPV16 is the most persistent type, but by 12 months 40% of infections clear or are treated because of diagnosis of a (pre)neoplastic lesion (Figure 3). This proportion reaches 85% by 36 months [3]. For HPV6, which is only very rarely associated with cancer [34] and instead most often with genital warts, these numbers are 66% and 98% respectively. These figures should nevertheless be taken with caution, as they directly depend on our ability to detect latent infections [18,24].

**Figure 3.**
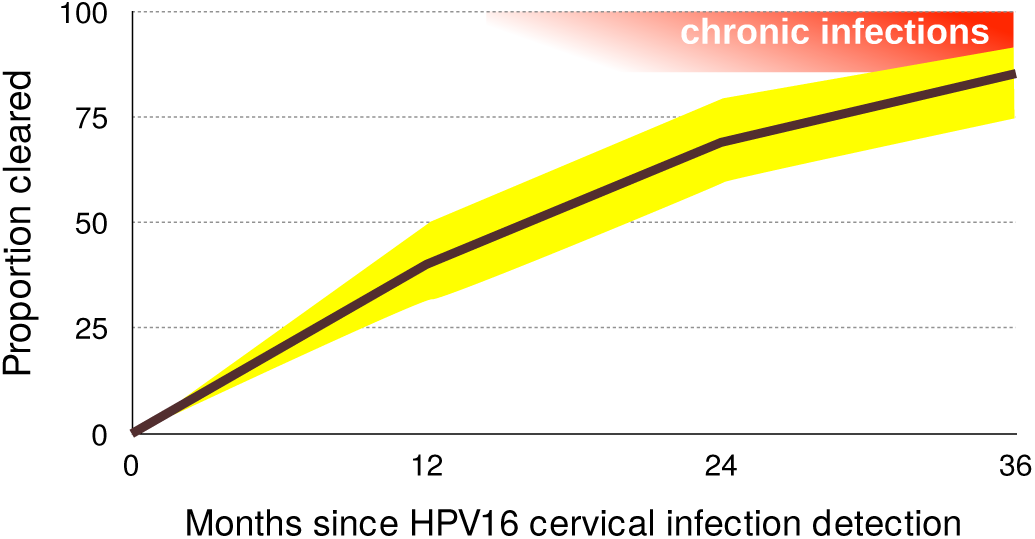
Proportion of HPV16 infections cleared 12, 24 and 36 months after the first detection. 95% confidence intervals are shown in yellow. Acute infections result in clearance or chronic infections, shown in red (note that chronic infections may clear as well). Data on clearance proportions originate from Table 4 in [3] and were obtained for 895 women in the United States of America, aged 16 to 23. Notice that chronic infections (in red) can also clear.

Besides studies on the natural history of the infection, the placebo arms of vaccine trials have been instrumental in increasing our understanding of the epidemiology of these infections by following thousands of young adults over time [e.g. 35–37]. In order to detect viral persistence of anogenital infections, the typical interval between two visits is usually six months. Studies with denser sampling (e.g. twice per week, as in [38]) often come at the expense of duration of the follow-up (16 weeks in this case), and therefore may provide accurate information about incidence but less information about persistence.

Some longitudinal studies performed more frequent sampling over a long period of time. For instance, two studies sampled women every 3 months. One of these followed unvaccinated girls aged 15-16 years in Tanzania and found that the median time from reported sexual debut to first HPV infection was five months, while median infection duration was six months [22]. Another study followed 150 adolescent women (aged 14 to 17) visiting primary care clinics for a median of six years [39], and found that 21% of HPV type-specific infections consisted of re-detection after apparent clearance (two negative visits). Note, however, that these girls were simultaneously infected by many HPVs, which could have generated some interference in the detection with the technique used. Furthermore, the population studied exhibited high rates of sexually transmitted infections (71.2% for chlamydia and 49.3% for gonorrhea) and a mean of above 10 sexual partners during the study. In spite of their limitations, these two studies suggest that the little we know about the duration and the actual clearance of acute infections could be challenged by studies with deeper resolution.

#### The immunology of clearance

Why most infections clear, while only few persist and progress to cancer remains largely unresolved (see Box 1 at the end of this article). Overall, we know that some cofactors such as HPV genotype, host genetic background, age of sexual debut, or coinfections, have an effect on clearance time [40]. However, our mechanistic understanding is still lagging behind.

The immunology of HPV infections has been thoroughly reviewed elsewhere [e.g. 41]. We know that HPV infections concur with a local anti-inflammatory environment, and that although the adaptive immune response is very efficient at clearing the infection, its activation is variable and sometimes insufficient to prevent future re-infections [36,41,42].

Recent research into the role of innate effectors (e.g. natural killer T cells) and Th-17 responses (e.g. *δ*_*γ*_ T cells) in cancerous lesions has provided new insights into chronic infections [43], but the implications for acute infection clearance is not obvious. More work, then, into innate and adaptive immunity activities during acute infections would help better elucidate mechanisms of clearance, as they are likely to be several. This is particularly true for non-cervical body sites, where the specific immunity microenvironment is less understood.

### Open challenges

## Deciphering HPV kinetics: the meta-population hypothesis

The extant genetic diversity of HPVs is enormous, with many different viral life styles. For anogenital infections alone, there is a large variance in infection duration, with values ranging from a few months [22] to years [3]. We propose here that the meta-population framework used in ecology can provide strong explanatory power and insight into this variance.

In ecology, a meta-population is a set of populations of the same species that are connected through dispersal [44,45]. Each population displays its own dynamics but the processes and patterns that emerge at a higher (meta-population) level can be very different from what happens within each population. This ecological framework has relevance for host-pathogen interactions. For instance, genetic data suggests that HIV infections may exhibit a meta-population structure in the spleen [46]. Also, within-host viral genetic structure such as this has been recently put forward as an underlying explanation for the great variance in set-point viral load observed between patients [47]. A meta-population framework was first proposed and applied to HPVs in a study of multiple type infections under natural and vaccinated immunological scenarios [48]. Here, we broaden this as a way to understand the variation in dynamics of HPV infections.

The modular nature of mammalian skin and the individual proliferation of cells set the scene for a meta-population scenario, one where discrete viral populations can establish in various ‘patches’ within the same anatomical ‘site’ and/or in various sites of the same host. We further argue that this framework can help explain the thin line between acute and chronic HPV infections.

Viral colonisation of an individual patch is largely a rare event, provided that the common assumption of access to basal cells through microlesions holds true. The ability of the virus to access other patches increases with infection duration, virion productivity and weakening of epithelium integrity. Following Ryser *et al.* [49], we argue that local extinction/success are stochastic processes. Some initially infected patches may not successfully infect a new patch before going extinct, while others might. To further complicate the picture, heterogeneity of cell types and skin structures leads to patch differences in susceptibility to infection, in permissivity to virion production, or in intensity of immune surveillance. Thus, certain patches will act as reservoirs (known as ‘source patches’ in ecology) while others will act as ‘sink patches’. In such source-sink dynamics, infection duration is driven by the extinction rate in the reservoir patches but the virus load detected depends on the number and productivity of the sink patches infected as well [47].

Regarding infection duration, the meta-population hypothesis allows to differentiate between genuine and apparent chronic infections. Genuinely chronic infections would refer to sustained and characteristic viral activity in a given patch for a long period of time, therefore allowing for the accumulation of molecular damage that may lead to malignisation [50,51]. Apparent chronic infections would be sequential transmission chains of infections from one patch to another within the same anatomical location (Figure 4). In this case, each individual patch infection may eventually die out, so that there is no sustained virus-cell interaction, even if the patient remains positive for the infection by the same viral type during a long time. Obviously, the repetitive infection events in these apparent chronic infections would increase the chances that one of them becomes genuinely chronic, if the virus-cell-environment interactions allow for it.

**Figure 4.**
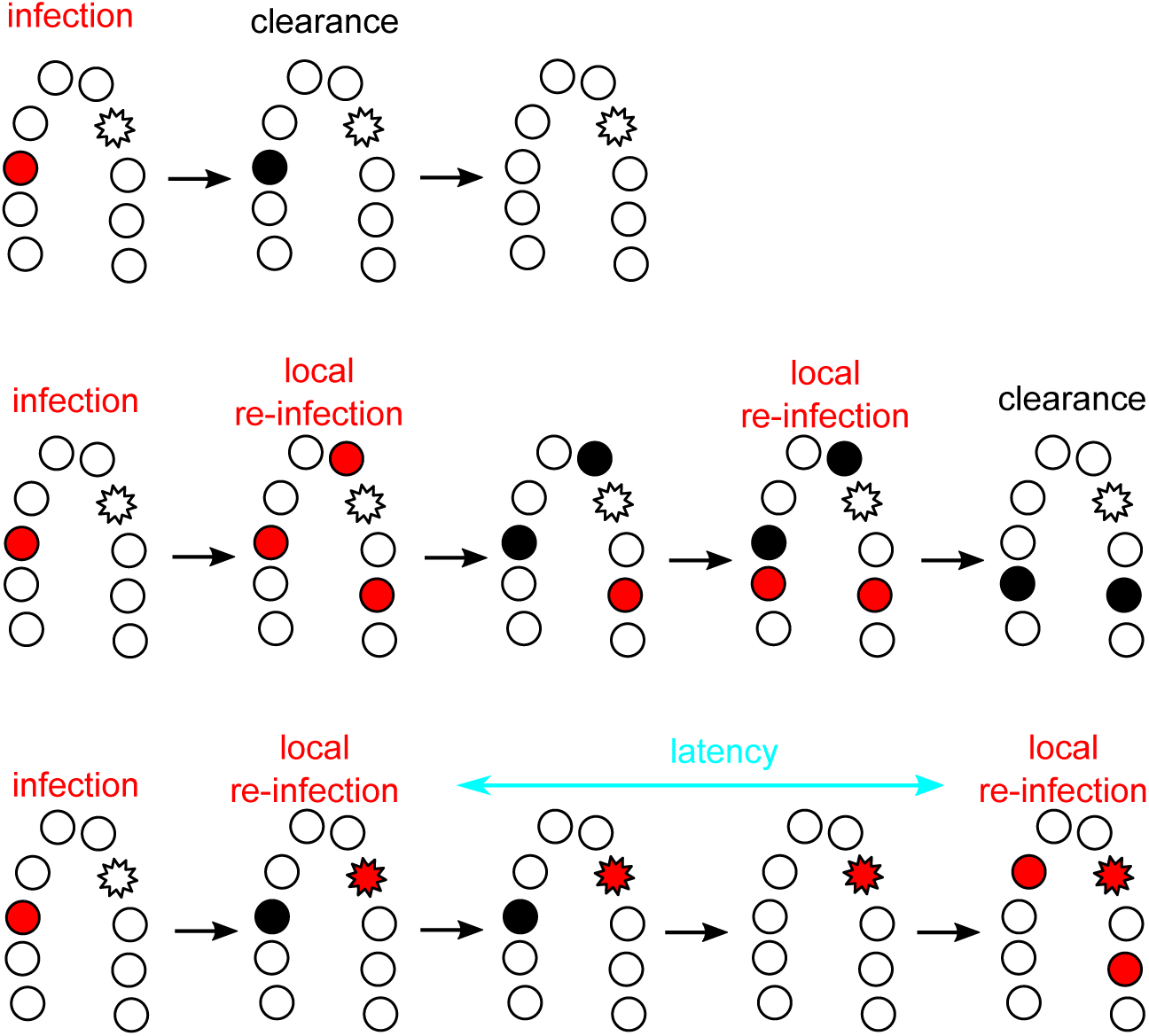
The meta-population hypothesis for HPV kinetics, exemplified in the cervix. Each circle represents a population (i.e. a cellular patch or a site) that can be susceptible (white), infected (red) or recovered from infection (black). The star-shape indicates a cellular patch that can remain infected for a longer time than the regular sites, and can that act as a source site if infected. For simplicity, recovered sites are here assumed to transitorily exhibit higher local immunity and resource depletion, thus making it unsustainable for re-infection. The top line shows a case with a local acute infection that is cleared without reaching other patches. The middle one shows a meta-population dynamics of local re-infection from an initial patch, without cell heterogeneity. The bottom line shows a case with patch heterogeneity (source-sink dynamics).

The meta-population framework, thus, may provide a powerful ecological explanation for the poorly understood differential progression of certain chronic infections towards cancer. Also, viral and cellular genomes will not accumulate similar genetic and/or epigenetic modifications during genuine and apparent chronic infections, thus allowing for molecular tests to differentiate between them. Testing the meta-population hypothesis will require HPV research to engage in spatial ecology studies, sampling between various macro/microscopic sites within the same individual (e.g. [52–55]), to study ‘auto-inoculation’ [56] and to systematically resort to microdissection studies to investigate the details of infected patches within sites [34,57–59].

## Cellular, viral and environmental heterogeneity: towards meta-communities

The mammalian skin is a complex environment with an intricate tri-dimensional structure. The most distinctive feature is the presence of hairs and the expression of keratins, keratin-associated proteins, the epidermal differentiation complex [60–62], as well as other hair evolutionary-related structures such as the mammary, sebaceous and sweat glands [63]. Different sites of the body have specific cell types, with unique composition, arrangement and density of skin appendages. Particular anatomical features are often observed at transitions between specialised epithelia, as in the tonsillar crypta, the transition zone in the cervix, the perianal region or the periungual area. Stratified skin renewal dynamics are strongly regulated and proceed vertically, with a gradient from the dividing cells in the basal layer to the differentiating and cornified keratinocytes in the outermost layer. The skin structure is, thus, conceivably modular, with columnar functional ‘patches’, which we will define here as the continuum of locations in the epithelium. The ‘vertical’ nature of the proliferating/desquamating keratinocyte patches is complemented by the ‘horizontal’ immune surveillance by Langerhans cells, dendritic cells and macrophages [64,65].

This heterogeneity goes beyond the strong assumption of the meta-population concept, which is that it only applies to populations, that is individuals from the same species. To account for different target cell types and for more ‘trophic interactions’ (here immune cells), we need to invoke the meta-community concept [66]. This has already proven to be appropriate to analyse within-host dynamics, because various parasite species often coinfect the same host, thus compete for space, energy, and resources [67,68].

A first advantage of an HPV meta-community approach is that it can factor in different target cell types. Papillomaviruses have been infecting amniotes since their first appearance [69] and have evolved strategies to successfully thrive across the diversity of structures and specialisations of present-day mammalian skin [70]. For instance, cells in hair follicles seem to act as reservoirs for chronic infection by cutaneous HPVs, while in stratified mucosal epithelia viral infections can become chronic in the basal cell layer. The columnar nature of the skin influences the natural history of the proliferative lesions induced by HPVs, which derive from the clonal expansion of basal/parabasal cells [70,71].

The heterogeneous nature of the skin underlies also the different propensity for chronic viral infections to lead to cancer in different anatomical locations. Many squamous cervical carcinomas associated with HPVs are thought to arise from a discrete population of cuboidal epithelial cells located in the transformation zone between the endo- and the ectocervix [72]. The absence of such transformation zones in the vagina, vulva or penis might explain why the cervix has a much higher burden of infection, higher incidence and younger age-at-diagnosis for cancers compared to other anatomical locations [73]. We can only speculate on why HPV-related cancers also occur in absence of a transformation zone, but a likely explanation is that all infected epithelial stem cells can potentially end up being carcinogenic but that the probability of such event depends on the cell phenotype and the local environment. Finally, heterogeneity between squamous and glandular patches results also in differential susceptibility towards chronification and malignisation by different HPVs and even by closely related viral variants, possibly also modulated by the host genetic background [74]. These known effects of cellular heterogeneity on chronic infections and cancer are likely to also affect acute infections but our knowledge about the latter is very limited.

More generally, the underlying hypothesis in our meta-population and meta-community approaches is that acute, latent and chronic infections exhibit differential specific features at the virocellular level, which further vary as a function of the infected cell type and differentiation status. Markers of acute *versus* true persisting infections will most likely be different for different anatomical sites and skin structures. The overall dynamics of the virus-host interaction is thus an integration of the interactions between viral genotype, host genotype, cellular phenotype and environment.

## HPV latency

Papillomavirus latent infections are still very poorly characterised and understood. The prevalence of latency remains largely unexplored, and its contribution to the oncogenic process in chronic infections is obscure. Establishment of latent infections is a life-history trait shared by very divergent viruses. For large DNA viruses, such as herpesviruses, the infection often starts with an acute phase, and then viral gene expression changes towards a different profile where the virus enters a latent state, with limited or even no genome replication, and no cellular damage, until reactivation is triggered. In small DNA viruses, such as Torque-Teno viruses, the initial infection goes unnoticed (the acute infection stage has actually never been documented) and the viral genomes remain in the host and replicate chronically at very low levels without apparent damage [2]. For papillomavirues, most of the evidence on latency originates from animal models, where latency is defined as low-level viral genome maintenance in the basal layers without a productive viral life cycle [25,75].

For HPVs, two ways for latency to arise have been proposed: infections may directly enter latency without going through an acute phase, or it may instead arise after a productive phase without successful clearance [24]. In both cases, latency is directly linked to acute infection dynamics. In the meta-communities context, latency can additionally be conceived to arise from heterogeneity in the interactions between virus and host cells. The molecular decision for cell division in the basal layer is stochastic and given that the viral genome requires cell division to replicate [76], latency-reactivation episodes could be mechanistically understood to reflect stochasticity in the time lapses of basal cell mitotic activity, without necessarily requiring viral manipulation of the host cell. Again, anatomical heterogeneity and viral genetic diversity may render certain anatomical locations combined with particular viral linages more prone to such latency-reactivation cycles. Overall, in addition to molecular virology investigations into latency mechanisms, studies combining a detailed initial follow-up (to demonstrate clearance), a long follow-up (to demonstrate chronic infection) and sequencing (to demonstrate that the virus causing the acute infection actually persisted) will help generate a complete picture on papillomavirus infection latency.

## Immunotherapies

Since vaccination will not be widespread for many years [77] and given that treatment interventions are often expensive and difficult to implement, the development of treatments remains urgent. Treatment development remains historically the less succesful front for HPV research. Understanding the mechanisms of HPV clearance promises aiding the development of immunotherapies, which consist in treating a disease by stimulating or suppressing the immune system.

Currently, the bulk of clinical and animal model research into HPV immunotherapies is understandably focused on cervical cancer treatment. Indeed, there are numerous approaches to developing these treatments, such as protein/peptide vaccines, bacteria-and-viral based vectors, and immunomodulators [78]. One of the most promising examples to date is the therapeutic vaccine VGX-3100, a plasmid containing synthetic versions of *E6* and *E7* genes of HPV16 and HPV18, which showed efficiency in a controlled trial at improving the regression of high-grade lesions (CIN2/3) [79]. The iatrogenic exposure to viral oncoproteins triggers a strong cellular immune response against the infected cells that the natural infection is not able to initiate. An alternative approach is to reverse the anti-inflammatory microenvironment that the virus creates during infections, in order to alert the immune system and boost its functioning [reviewed in 80]. While several of these therapies have reached clinical trial stages, it has become clear that one mechanism alone (e.g. augmenting CTL infiltration and function) will not suffice, and combinations of mechanisms are needed. Studying how these mechanisms work in natural acute infections might help.

In practice, immunotherapies cannot be envisaged to treat all acute HPVs infections given their prevalence and their often sub-clinical presentations. However, knowledge onthe acute stage could be transferable to fight chronic stage infections. For instance by identifying any immune stimulants or immune cell subtypes that help clear natural acute infections. However, it may be possible that several treatments will be needed for different lesion grades (e.g. for low to high-grade neoplastic lesions than for late-stage cancers), given that the heightened immune suppression microenvironment in advanced malignancies is particularly complex and strong. Therefore, insights from acute infection clearance could be particularly well suited for development of therapies against premalignant lesions that are usually removed surgically.

More focused applications could arise from studying HPV infections in sites other than the cervix, for instance in the case of respiratory recurrent papillomatosis. This is a rare condition caused by chronic infection by HPV6 or HPV11 that shares features of both acute and chronic infections. The chronic disease imposes a recurrent burden to the patients to control the acute, benign clinical presentations of the disease, which can progress fatally to the lungs [81]. Furthermore, given the unique microenvironments of non-cervical sites, studies of infection dynamics at these sites are greatly needed. In particular, anal intraepithelial neoplasias and anal squamous cell carcinomas are increasing in prevalence worldwide yet studies of HPV-immunity interactions in anal infections are few [82].

## Fertility

The Zika epidemic has reminded us of the risk viral infections have on pregnancies [83]. Anogenital infections by HPVs deserve to be studied in this context because rare deleterious effects could translate into an important burden, given their high prevalence. There is a well-established link between cervical disease and pregnancy complications [84], but we are only beginning to understand the connection between clinically asymptomatic HPVs infections and complications such as pre-eclampsia, fetal growth restriction or pre-term delivery [85].

In men, anogenital asymptomatic infections by mucosal and cutaneous HPVs are very common [37] and some are associated with penile cancer [5]. Surprisingly, HPV DNA can also be found in human semen [86] and although this viral DNA could originate from desquamating epithelial cells, data suggest the presence of viral DNA directly associated to sperm cells [23]. Evidence supporting a correlation between infection by HPVs in men and infertility is still controversial [87] and the mechanisms involved remain largely unknown [88].

Finally, *in utero* transmission of HPVs is rare [89], but asymptomatic cervical viral infections can reduce barrier integrity [90] and viral DNA can be detected in the placentae [91]. There is further a sound body of evidence describing transmission of HPVs during vaginal birth, as illustrated by the case of infantile respiratory recurrent papillomatosis described above [81].

Overall, the presence of mucosal HPVs in placenta and semen suggests non-classical tropisms, and viral life cycle in these particular cellular environments needs to be elucidated. Clinical studies investigating acute anogenital infections in pregnant women (along the various stages of pregnancy) as well as in their partners are required to unravel how HPVs (independently of their oncogenic potential) may be either directly or indirectly increasing the risk of infertility, spontaneous abortions, pre-term labor, preeclampsia or other complications.

## Vaccine escape

Effective vaccines with high coverage rates exert major selective pressures on pathogens [92]. The risk of an evolutionary response from the pathogen relies on its genetic diversity, and does not necessarily require *de novo* mutations if it is already diverse. Extant diversity in papillomaviruses is immense, with hundreds of viral lineages, which themselves harbour significant genetic variation [93].

While vaccines against HPVs are effective and safe, they may be leaving the door open for an evolutionary response [48]. This can occur if some vaccinated individuals are infected by HPV vaccine types (due to infection before vaccination, immunosuppression or failure to mount a sufficiently strong immune response). In the clinical trial for the nonavalent vaccine, even in the per-protocol efficacy population, where the *n* = 5812 participants received all three doses of vaccine within one year and were HPV-uninfected at inclusion, the median number of infections by one of the targeted viruses was 3.6 for 1,000 person-years at risk (data from Table S5 in [94]). In the intention-to-treat population, which did not exclude 988 additional participants who received at least one vaccine dose, this number increased to 36. Despite the outstanding vaccine efficacy, when millions of women are vaccinated, this will still represent thousands of infections by vaccine-targetted viral genotypes that would be eventually cleared and not result in malignant disease. These acute infections would thus occur in vaccinated women, with a strong immune response, and the question arises whether they may last long enough to allow specific viral lineages (either pre-existing or *de novo* generated viral variants) to be differentially transmitted, thus paving the way for viral adaptation to this special environment.

HPV vaccines can generate off-target immune activity and offer protection against closely related viruses not targeted by the vaccine formulation [95,96]. Yet viral diversity remains so large that in the few regions where vaccination has reduced the prevalence of vaccine-targeted types, most other HPVs continue to circulate [6,97]. Overall, vaccination against HPVs will undoubtedly have a strong and highly desirable impact in disease prevention, but it will create novel host environments to which viruses may adapt.

Thanks to next generation sequencing, we now have access to an increasing number of full genomes. For instance, very recently, an analysis of more than 5500 HPV16 genomes identified shared feature between viruses isolated from precancers and cancers compared to viruses from case-control samples [98]. However, evaluating the potential for HPV evolution will require genomic data of intra- and inter-individual viral populations collected through time. Since HPVs are double-stranded DNA viruses replicated by host polymerases, mutation rates are expected to be low. Nevertheless, we know little about polymerase fidelity in somatic cells, both in general and during a viral infections. Investigation of long follow-up data sets with short sampling intervals are lacking, especially since evolutionary rates may exhibit periods of rapid increase followed by long periods of stasis, as demonstrated for Influenza A virus [99]. Further, in the case of HPVs, displaying acute, chronic and latent stages of the infection, the evolutionary dynamics may strongly depend on the presence of a latent phase and on the epidemiological transmission patterns, as described for Varicella-zoster virus [100]. For HPVs, focusing on the acute infections could therefore yield novel insights since this is where the viral life cycle is most productive.

## Scars that matter long after clearance?

Even when viral clearance does occur, recent work shows that acute infections can impair the immune system causing chronic inflammation or ‘immunological scaring’ [105]. Certain oncoviruses are believed to leave behind molecular damage in their host cells, which can lead to cancer several years later [106]. Cervical cancers are a clear example of direct carcinogenesis, since chronic infection by oncogenic HPVs is a necessary cause for virtually all of these cancers. However, even acute infections can induce modifications in the cellular (epi)genome, creating the stage for pre-cancerous lesions [106,107]. Although an old hypothesis, the ‘hit-and-run’ effects of acute infections are poorly understood for bacterial or viral infections, and exploring how HPVs may cause this kind of damage remains an important research direction [108].

Speculatively, such a long-term impact could concern not only anogenital but also infections at cutaneous sites. Indeed, virtually all humans become infected by very diverse cutaneous HPVs, chiefly beta- and gammapapillomaviruses, which are proposed to act as cofactors for the risk of developing non-melanoma skin cancers in certain human populations [109]. Given that such HPVs infections are extremely prevalent (Figure 5), often causing subclinical infections and interact extensively with the immune system, they could be an ideal model for studying ‘under-the-radar’ viral infections and their potential side-effects on immune functioning.

**Figure 5.**
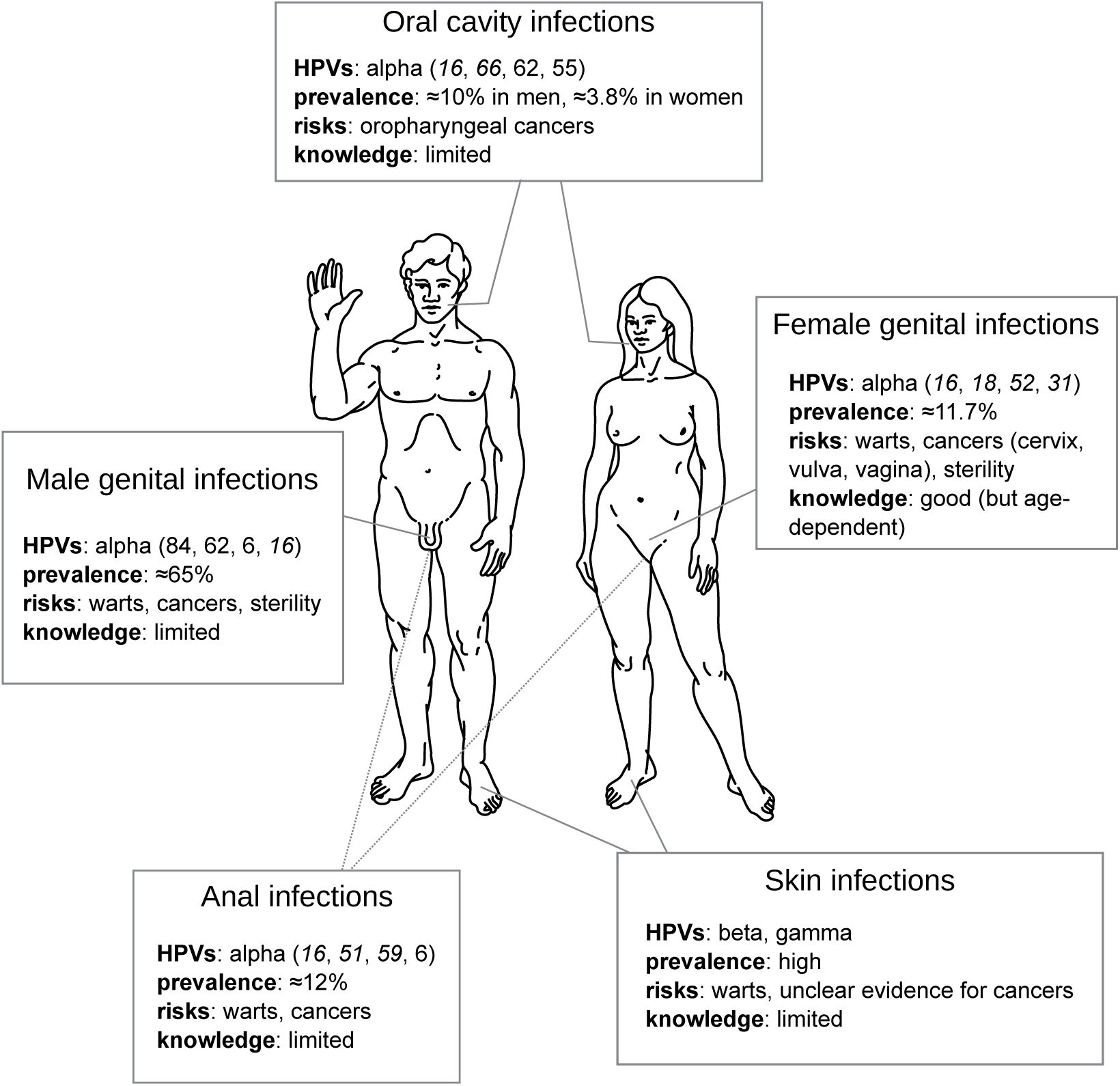
Challenges of HPV acute infections per anatomical location. For each location, HPVs are ordered per prevalence and oncogenic ones are in italic. Most prevalences are global estimates [1,4,101], except for oral (US [102]), genital in men (Brazil, Mexico, US, [103]) and anal (Brazil, Mexico, US [104]) sites. Detailed data are available in Supplementary Table.

## Conclusions

Acute infections by HPVs are the rule rather than the exception, and a better understanding of such infections is urgent because of their enormous fundamental and public health implications. We propose that metapopulation and metacommunity approaches borrowed from ecology can provide a strong explanatory framework to the study of the course of HPV infections and better distinguish between acute, latent and chronic infections. In addition to the promise of identifying early markers of infection chronicity, understanding viral-immunity interactions can help design new treatments to boost natural immunity against novel infections. Such therapies may reduce the chances of an acute infection to chronify and to reach (pre)cancer stages. Detailed studies with long follow-up and short time intervals are further needed to precisely assess the role acute HPV infections could have on fertility or on the long-term ‘immune scars’ these infections may leave behind.

Finally, as with most infectious diseases, it is important to remain humble and to accept that elimination [110] of virulent HPVs is unlikely. In fact, it is not clear whether such selective removal should be desirable, as most HPVs are part of our virome and might be playing important ecological roles. This is why the development of vaccines should not stop us from improving our characterization of the natural history of HPVs and to shift from an elimination to control perspective.

#### Box 1: Open questions about HPV acute infections

- Do some innate immunity evasion mechanisms lead to delayed clearance? [43]
- How common is immunological tolerance to HPVs?
- What role, if any, do natural B-cell responses play in clearance?
- Are there differences in immunological clearance mechanisms for different HPVs or for different infected cell types, tissue structures or anatomical sites?
- Are there long term effects after HPV infection clearance (immunological scaring)? • How do HPV infections affect male and female fertility?
- How do HPV infection kinetics affect transmission? [111]
- Are HPVs infections structured in host tissues (the meta-population hypothesis)?
- What is the role of the microbiota in HPV infection clearance, persistence and progression to cancer? [112,113]

## Supplementary Materials

The following are available online at <u>www.mdpi.com/link</u>, Table S1: HPV epidemiological data for Figure 5.

## Acknowledgments

SA is supported by the European Research Council (ERC) under the European Union’s Horizon 2020 research and innovation program (EVOLPROOF, grant agreement No 648963). NB is supported by the European Research Council (ERC) under the European Union’s Horizon 2020 research and innovation program (CODOVIREVOL, grant agreement No 647916). All authors acknowledge further support from the CNRS and the IRD.

## Author Contributions

All authors conceived and wrote the review.

## Conflicts of Interest

The authors declare no conflict of interest. The funding sponsors had no role in the writing of the manuscript, and in the decision to publish it.

## Abbreviations

The following abbreviations are used in this manuscript:

CIN: Cervical Intraepithelial Neoplasia
CTL: Cytotoxic T Lymphocytes
HPVs: Human Papillomaviruses
HR: High Risk
LR: Low Risk
UN: United Nations

